# Characterization of redox sensitive algal mannitol-1-phosphatases of the haloacid dehalogenase superfamily of proteins

**DOI:** 10.1101/2020.07.01.179531

**Authors:** Yoran Le Strat, Thierry Tonon, Catherine Leblanc, Agnès Groisillier

## Abstract

Macroalgae (or seaweeds) are the dominant primary producers in marine vegetated coastal habitats and largely contribute to global ocean carbon fluxes. They also represent attractive renewable production platforms for biofuels, food, feed, and bioactives, notably due to their diverse and peculiar polysaccharides and carbohydrates. Among seaweeds, brown algae produce alginates and sulfated fucans as constituents of their cell wall, and the photoassimilates laminarin and mannitol for carbon storage. Availability of brown algal genomes, including those of the kelp *Saccharina japonica* and the filamentous *Ectocarpus* sp., has paved the way for biochemical characterization of recombinant enzymes involved in their polysaccharide and carbohydrates synthesis, notably mannitol. Biosynthesis of mannitol in brown algae starts from fructose-6-phospate, which is converted into mannitol-1-phosphate (M1P), and this intermediate is then hydrolysed by a haloacid dehalogenase type M1P phosphatase (M1Pase) to produce mannitol. We report here the biochemical characterization of a second M1Pase in *Ectocarpus* sp after heterologous expression in *Escherichia coli*. (EsM1Pase1). Our results show that both *Ectocarpus* M1Pases were redox sensitive, with EsM1Pase1 being active only in presence of reducing agent. Such catalytic properties have not been observed for any of the M1Pase characterized so far. EsM1Pases were specific to mannitol, in contrast to *S. japonica* M1Pases that can use other phosphorylated sugars as substrates. Finally, brown algal M1Pases grouped into two well-supported clades, with potential different subcellular localization and physiological role(s) under diverse environmental conditions and/or stages of life cycle.

## 1. Introduction

Macroalgae, or seaweeds, are the dominant primary producers in marine vegetated coastal habitats and largely contribute to global ocean carbon fluxes (Krause-Jensen et al., 2018). Seaweeds represent also attractive bioresources for biofuels, nutraceuticals or therapeutics (Kawai and Murata, 2016; Abdul Khalil et al., 2017; Sudhakar et al., 2018). Based on a recent analysis (https://www.alliedmarketresearch.com/seaweed-market), the global seaweed market is expected to reach $9.1 billion by 2024, driven by the growing application of macroalgae in various industries. One aspect making macroalgae attractive to industry is their high, diverse, and peculiar polysaccharide and carbohydrate contents compared to terrestrial plants (Wei et al., 2013; Rioux and Turgeon, 2015). Seaweeds are part of a polyphyletic group including green algae, red algae, and brown algae. These latter contain in their wall the complex polysaccharide alginates and sulphated fucans, and store photosynthesis-derived carbon by producing the beta-1,3-glucan laminarin and the sugar alcohol mannitol. This polyol can represent between 20 to 30% of the dry weigh of brown seaweed (Reed et al., 1985).

Recent advances in the molecular bases of biosynthetic pathways of cell wall polysaccharides (alginates, sulfated fucans) and mannitol have been made in *Saccharina japonica* and *Ectocarpus* sp., notably through heterologous expression of candidate genes identified in their respective genomes (Ye et al., 2015; Cock et al., 2010). Enzymatic characterization of recombinant proteins confirmed the functions of genes coding for proteins involved in mannuronan and fucoidan metabolisms of *Ectocarpus* sp. and of *S. japonica* (Tenhaken et al., 2011; Zhang et al., 2016; Chi et al., 2018a). It also enabled the biochemical analysis of several C5-epimerases catalyzing the last step of alginate production (Fischl et al, Glycobiology 2016; Inoue et al., 2016), and more recently of the first algal alginate lyase (Inoue and Ojima, 2019). Despite progresses in the recent years, heterologous expression of brown algal genes and subsequent purification of recombinant proteins remain challenging, with most of the successful expression obtained so far using *E. coli* as a host (Groisillier, 2018).

Regarding mannitol metabolism, changes in the content of this polyol in *Ectocarpus* sp. follow a diurnal cycle (Gravot et al., 2011), with higher quantities at the end of the light period. Yields of mannitol are also influenced by seasonal variations, with higher amounts during summer and autumn months (Schiener et al. 2015). Mannitol biosynthetic pathway in brown algae relies on two enzymatic activities: mannitol-1-phosphate dehydrogenase (M1PDH) converts fructose-6-phosphate (fructose-6P) into mannitol-1- phosphate (mannitol-1P), this intermediate being further transformed into mannitol by haloacid dehalogenase (HAD) type mannitol-1P phosphatase (or mannitol-1- phosphatase, M1Pase). Potential genes coding for such enzymes have been identified in *Ectocarpus* sp. by mining genomic resource, i.e. three candidates for M1PDH and two for M1Pase (Michel et al., 2010). In this alga, one M1PDH (Rousvoal et al., 2011; Bonin et al., 2015) and one M1Pase (EsM1Pase2, Groisillier et al., 2014) have been previously biochemically characterized by heterologous expression in *Escherichia coli*. Very recently, Chi et al. (2018b) have identified and characterized homolog genes in *S. japonica*. Other M1Pases have been characterized from diverse organisms. Native enzymes have been purified from the red macroalga *Caloglossa continua* (Iwamoto et al., 2001), or partially purified from the brown macroalgae *Spatoglossum pacificum* and *Dictyota dichotoma* (Ikawa et al., 1972). Recombinant proteins have been studied for the phosphohistidine phosphotransferase M1Pase of the chicken parasite *Eimeiria tenella* (Liberator et al., 1998), and of the HAD M1Pase module of a M1Pase/M1PDH fusion from the soil bacteria *Acinetobacter baylyi* (Sand et al., 2015). A fusion protein of the green alga *Micromonas pusilla* containing a M1PDH module fused in C-terminal with a HAD type M1Pase module was shown to enable production of mannitol in both recombinant *E. coli* and cyanobacteria (Madsen et al., 2018). There are important differences between both *Ectocarpus* sp. M1Pases (EsM1Pases). Compared to EsM1Pase2, EsM1Pase1 features an unique N-terminal extension (85 aa) of unknown function, and has been predicted to localize in the chloroplast. Previous attempts to purify recombinant native full-length and truncated (i.e. without the 85 aa extension) EsM1Pase1 have failed (Groisillier et al. 2014). Here we report the successful characterization of a codon-optimized EsM1Pase1 gene in *E. coli* after deletion of its signal peptide. We observed that both *Ectocarpus* sp. M1Pases were specific to mannitol, in contrast to results obtained for orthologs in *S. japonica*. In addition, we observed that both EsM1Pases were redox sensitive, with EsM1Pase1 being active only in presence of reducing agents. Finally, brown algal M1Pases grouped into two distinct clades that may have evolved to support mannitol production in different cellular compartments, and under different environmental conditions and/or during different stages of life cycle.

## 2. Results and discussion

### 2.1 EsM1Pase1 is a bona fide M1Pase

We have previously reported attempts to purify recombinant native full-length EsM1Pase1, as well as of a native truncated form in which the entire N-terminal extension (255 nt et al., 85 aa) was removed (Groisillier et al., 2014). Despite considering different expression plasmids, host cells, and induction conditions, no soluble protein was produced in sufficient quantity.

Wild type *Escherichia coli* is not able to produce mannitol and does not contain any M1Pase gene. To assess if EsM1Pase1 corresponds to a genuine M1Pase, *E. coli* cells were transformed with plasmids containing full-length native or full-length codon-optimized EsM1Pase1. *E. coli* containing a plasmid with the gene coding for EsM1Pase2 was used as a positive control (Groisillier et al., 2014). Recombinant *E. coli* cells were tested for their capability to produce mannitol in minimal medium containing glucose. Figure 1 showed that both native and codon-optimized EsM1Pase1 were functionally expressed in *E. coli*, as they triggered the production of mannitol only in presence of IPTG. Similar levels of mannitol were measured in the culture medium for the three proteins tested, ranging between 0.276 ± 0.026 to 0.319 ± 0.014 g/l. These results supported the prediction that EsM1Pase1 was an effectiveM1Pase, and paved the way for subsequent biochemical characterization of the recombinant codon-optimized EsM1Pase1.

**Fig. 1.**
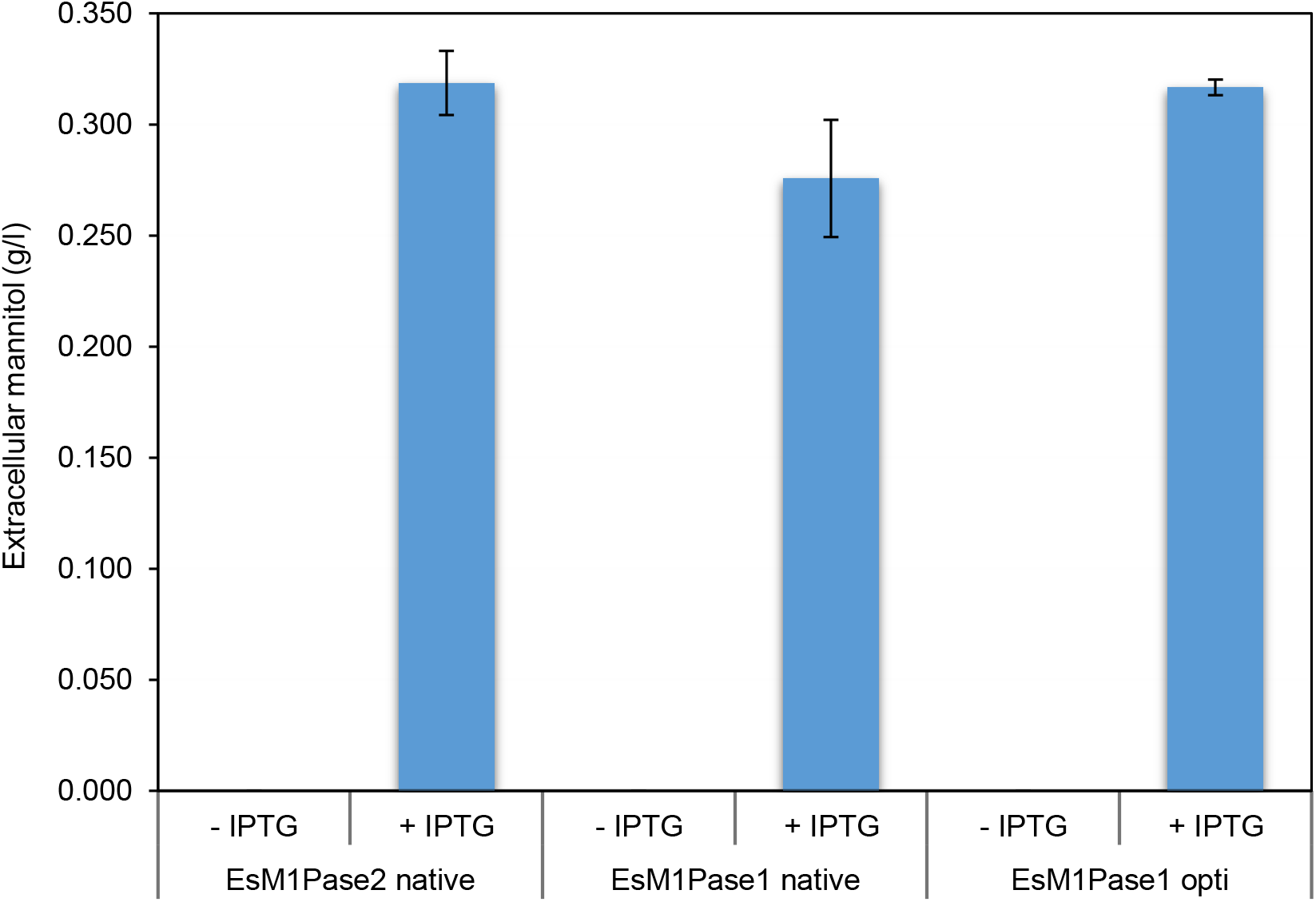
Mannitol production after cultures of recombinant *E. coli* expressing *EsM1Pase* genes. Mannitol concentration in the media of *E. coli* cultures transformed with *EsM1Pase1* or *EsM1Pase2* genes under the control of T7 promoter was determined in presence and in absence of IPTG. No mannitol was detected in absence of induction by IPTG. Data presented are means ± S.D. from culture of three independent clones for each gene tested. opti, full-length codon-optimized sequence for expression in *E. coli*.

### 2.2 Biochemical characterization of EsM1Pase1short recombinant proteins

To improve recombinant expression and further purification of EsM1Pase1, a truncated version of the codon-optimized gene, named *EsM1Paseshort*, was cloned in the plasmid pFO4. The deleted sequence corresponded to the first 117 nucleotides of the gene, and coded for a potential plastid signal peptide of 39 aa. After cloning and transformation of *E. coli*, the production of soluble recombinant proteins was obtained under double induction by lactose and IPTG in LB medium. Recombinant His-tagged EsM1Pase1short proteins were purified to homogeneity by a two-step protocol based on Ni^2+^-affinity chromatography (Fig. 2A) and gel filtration (Fig. 2B). Presence of EsM1Pase1short protein (theoretical mass of 41 kDa) in collected fractions was confirmed by SDS-Page and Western-Blott using anti-histidine tag antibodies (Fig. 2C and 2D respectively), and by measuring M1P phosphatase activity (data not showed). Gel filtration profile showed two peaks. The first corresponded to inactive EsM1Pase1short aggregates, and the second to the active form of the enzyme of interest. Estimation of molecular mass of proteins (around 40 kDa) contained in the second peak indicated that EsM1Pase1short was functional as a monomeric form in solution. In algae, only two other quaternary structure of M1Pase have been determined so far, *i.e.* for the enzyme of the red alga *Caloglossa continua* (Iwamoto et al., 2001), active as a monomer, and for the EsM1Pase2 from *Ectocarpus* sp. that is active as a tetramer (Groisillier et al., 2014). The two HAD type M1Pases from *Saccharina japonica* (SjaM1Pases) have also been purified to homogeneity (Chi et al., 2018b), but no information related to quaternary structure were given.

**Fig. 2.**
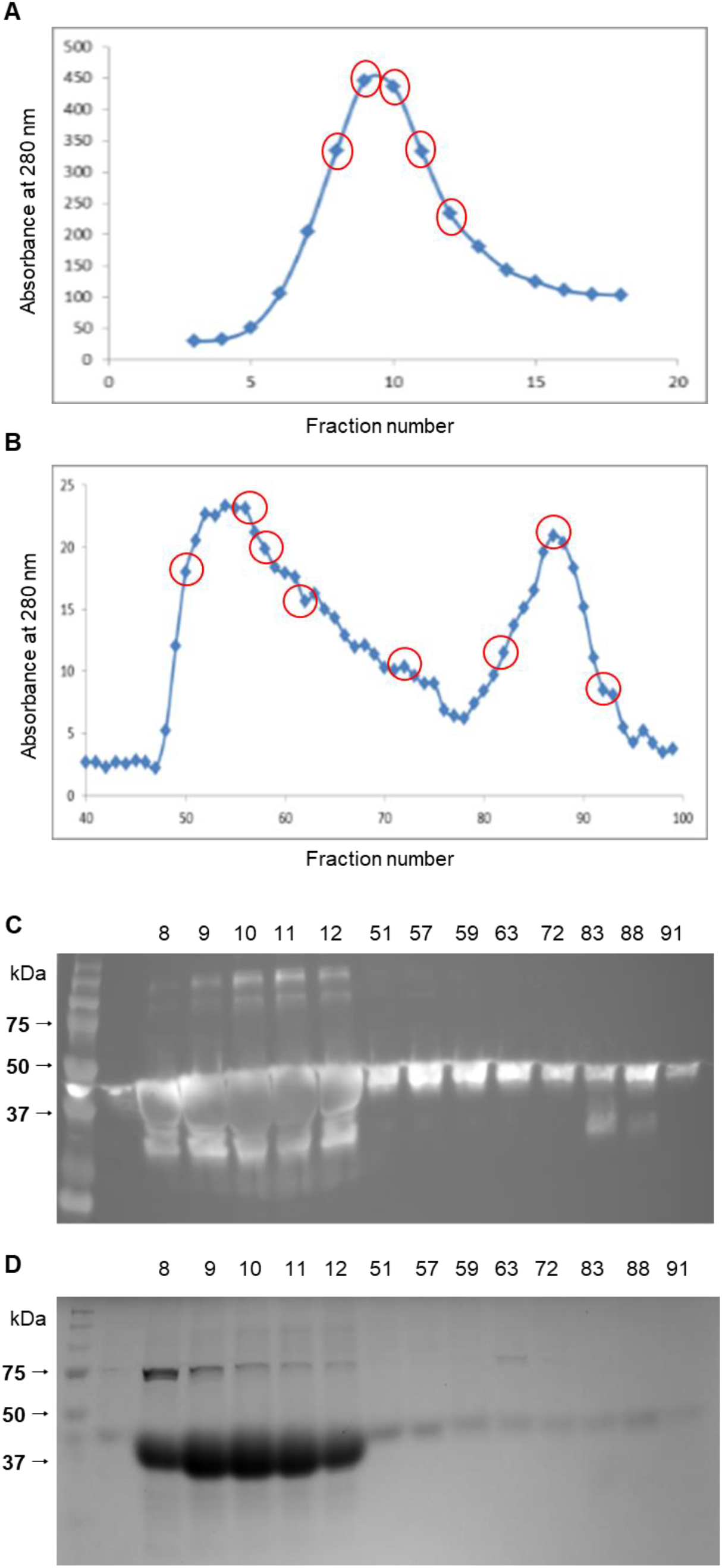
Purification of the recombinant His-tagged EsM1Pase1short. Proteins were purified first by Ni^2+^ affinity chromatography (A), and then were resolved by gel filtration onto a Superdex 200 HiLoadTM column (B). Red circles correspond to fractions from affinity purification (8 to 12), and from gel filtration (51 to 91), deposited for SDS-PAGE (C), and Western-Blot (D) analysis.

Preliminary activity tests using purified EsM1Pase1short (fractions 83 to 91 of Fig. 2B) were performed in presence of 1 mM mannitol-1P et al., 100 mM Tris-HCl pH 7.5, and 5 mM MgCl_2_ (final concentration) as described previously for EsM1Pase2 (Groisillier and Tonon, 2016), but no M1Pase activity was detected. Interestingly, during our previous study on recombinant EsM1Pase2, we observed a very low stability of this protein illustrated by a loss of activity less than 20 hours after purification. In addition, recent results have shown the redox sensitivity of a mammalian HAD-type phosphoglycolate phosphatase: its activity was inhibited by oxidation, but could be re-activated by reduction (Siefred et al et al., 2016). In this context, enzymatic activity of EsM1Pase1short was tested in presence of the reductant dithiothreitol (DTT) at 3 mM final concentration. Under these reducing conditions, the enzyme was found to be active. In Groisillier et al. (2014), the kinetic analyses of EsM1Pase2 were done just after the purification because the enzyme was not stable in the conditions tested. Based on the positive effect of DTT on recombinant EsM1Pase1short, EsM2Pase activity was also tested in presence of 3 mM DTT. Presence of the reductant in the assay mixture permitted to maintain enzymatic activity of the EsM1Pase2 tetramer for at least five days after purification (data not shown). Interestingly, none of the purified native and recombinant M1Pases biochemically characterized so far have been shown to be redox sensitive (Ikawa et al., 1972; Liberator et al., 1998; Iwamoto et al., 2001; Sand et al., 2015; Chi et al., 2018b).

Based on these results, all the following analyses for EsM1Pase1short were achieved in presence of 3 mM DTT. The specificity of EsM1Pase1short was determined by assaying activity in presence of different potential substrates at 1 mM final concentration. As no activity was detected with other substrates, this enzyme was found to be specific to mannitol-1P (Table 1), as also observed for EsM1Pase2 (Groisillier et al., 2014). Such narrow substrate specificity has been previously observed for M1Pases characterized in the brown algae *Dictyota dichotoma* and *Spatoglossum pacificum* (Ikawa et al., 1972), and in the red alga *Caloglossa continua* (Iwamoto et al., 2001). In contrast, in the brown alga *S. japonica*, significant phosphatase activity for both SjaM1Pases was also detected in presence of others substrates, such as glucose-1P, glucose-6P, and fructose-6P. Furthermore, the highest activity of SjaM1Pase2 was measured with glucose-1P (Table 1) (Chi et al., 2018b), as also observed for the M1Pase of the red alga *Dixoniella grisea* (Eggert et al., 2006). However, this latter result should be taken with caution because activities were measured on algal crude extracts, and not with (partially) purified proteins.

**Table 1.** Comparison of substrate specificity of biochemically characterized brown algal M1Pases. Results are expressed in percentage of activity using the activity measured in presence of mannitol-1P as 100 %, except for SjaM1Pase2.

|  | Mannitol-1-P | Glucose-1-P | Glucose-6-P | Fructose-6-P | Fructose-1-P | Mannose-6-P |
| --- | --- | --- | --- | --- | --- | --- |
| *S. japonica*<br>(SjaM1Pase1)<sup>1</sup> | 100 | 27.5 | 21.3 | 8.7 | n.d | n.d. |
| *S. japonica*<br>(SjaM1Pase2)<sup>1</sup> | 89.8 | 100 | 69.6 | 34.8 | n.d. | n.d. |
| *Ectocarpus sp.*<br>(EsM1Pase1short)<sup>2</sup> | 100 | 0 | 0 | 0 | 0 | 0 |
| *Ectocarpus sp.*<br>(EsM1Pase2)<sup>3</sup> | 100 | 0 | 0 | 0 | 0 | 0 |

n.d., not determined.
<sup>1</sup>, Chi et al., 2018b.
<sup>2</sup>, This study.
<sup>3</sup>, Groisillier et al., 2014.

The purified EsM1Pase1short protein exhibited a typical Michaelis-Menten kinetic when assayed with M1P concentration ranging from 0.0625 mM to 1.25 mM. Apparent K_m_ and V_m_ were determined from the Lineweaver-Burk plots (Fig. S1). For a better comparison between EsM1Pase1short and EsM1Pase2, the K_m_ and V_m_ of the latter enzyme was determined in presence of 3 mM DTT (Table 2, Fig. S2). Following the addition of the redox agent, the V_m_ value of EsM1Pase2 was multiplied by 2.7, the K_m_ doubled, and the K_cat_ of the protein increased from 0.02 to 0.05 s^−1^. However, EsM1Pase1short is much more active than EsM1Pase2. Indeed, specific activity and K_cat_ of EsM1Pase1short are 15 and 16 times higher than those of EsM1Pase2, respectively. As indicated in Table 2, similar biochemical properties were observed when comparing kinetic constants of SjaM1Pase1 and SjaM1Pase2 on mannitol-1P. In *S. japonica*, it was suggested, based on gene expression analysis, that SjaM1Pase1 was the main M1Pase responsible for production of mannitol, whereas SjaM1Pase2 may support mannitol synthesis under changes in environmental conditions (Chi et al., 2018b). Interestingly, in *Ectocarpus* sp., the gene coding for EsM1Pase2 was not found to be differentially expressed under short-term abiotic stress conditions, while *EsM1Pase1* was down-regulated under oxidative and hyposaline conditions (Dittami et al., 2009).

**Table 2.**
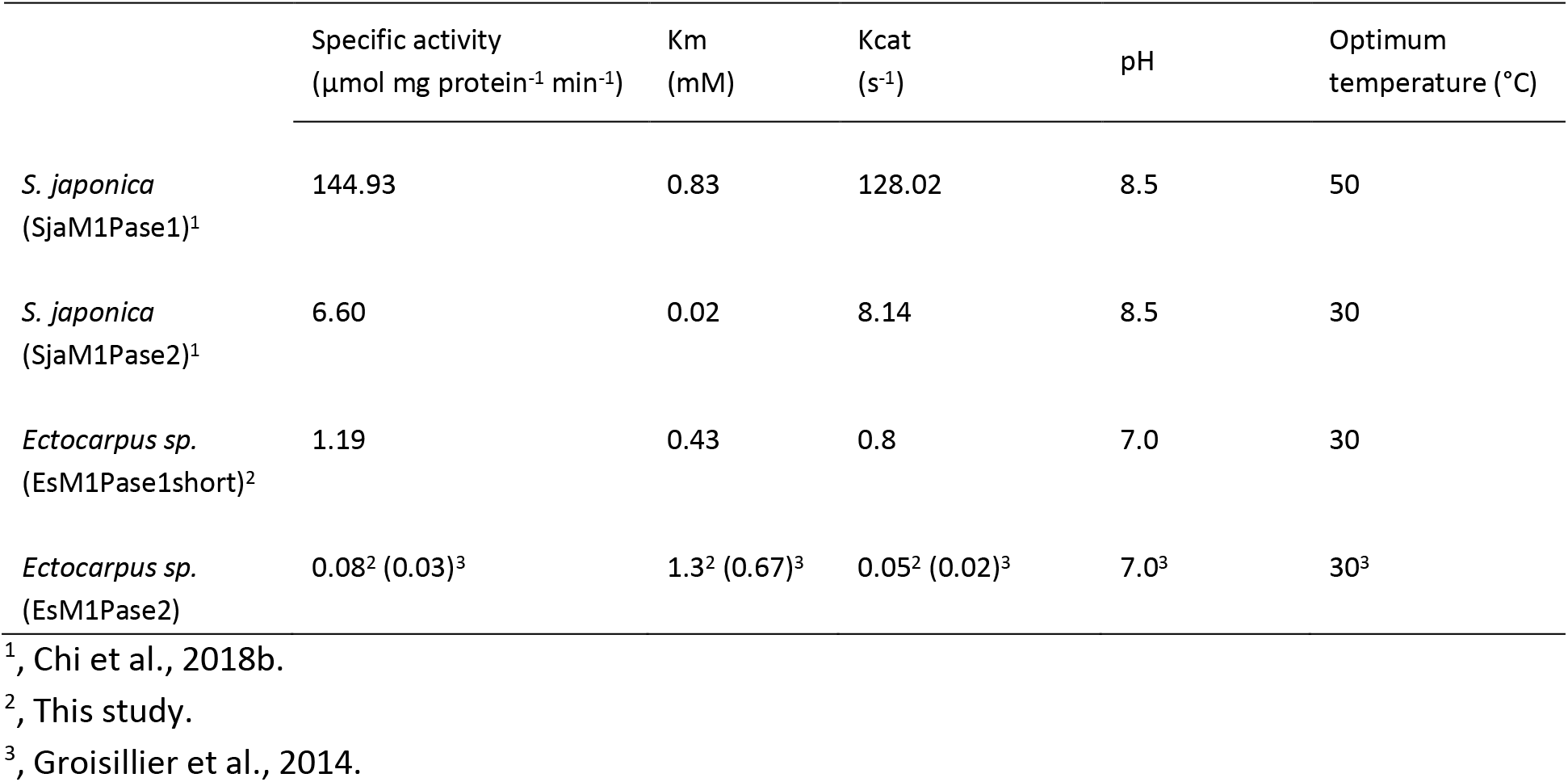
Comparison of kinetic properties of biochemically characterized brown algal M1Pases.

The optimum pH for EsM1Pase1short activity was 7.0, with 61 % and 83 % of the maximum activity remaining at pH 6.5 and pH 8.0 in 0.1 M Tris-HCl buffer, respectively (Fig. 3A). This is in agreement with pH values found in different brown and red algae, except for *S. japonica* whose optimum pH was 8.5 for both SjaM1Pases. The highest enzyme activity was observed at 30 °C in Tris-HCl, pH 7.0. The enzyme was still active between 4 and 12 °C (about 30 % of activity), while activity dropped to less than 8 % at 50°C (Fig. 3B). For comparison, the optimum temperature was also 30° C for EsM1Pase2 (Groisillier et al., 2014) and for SjaM1Pase2, but was found to be 50 °C for SjaM1Pase1 (Chi et al., 2018b). The activity of recombinant EsM1Pase1short significantly decreased with increasing NaCl concentration (Fig. 3C). About 30% of the initial activity was measured in presence of 1 M NaCl, while it was only 15 % for EsM1Pase2 (Groisillier et al., 2014) and 60 % for both *S japonica* enzymes (Chi et al., 2018b). This suggests that SjaM1Pases may be more tolerant to high NaCl concentrations than their counterparts in *Ectocarpus* sp.

**Fig. 3.**
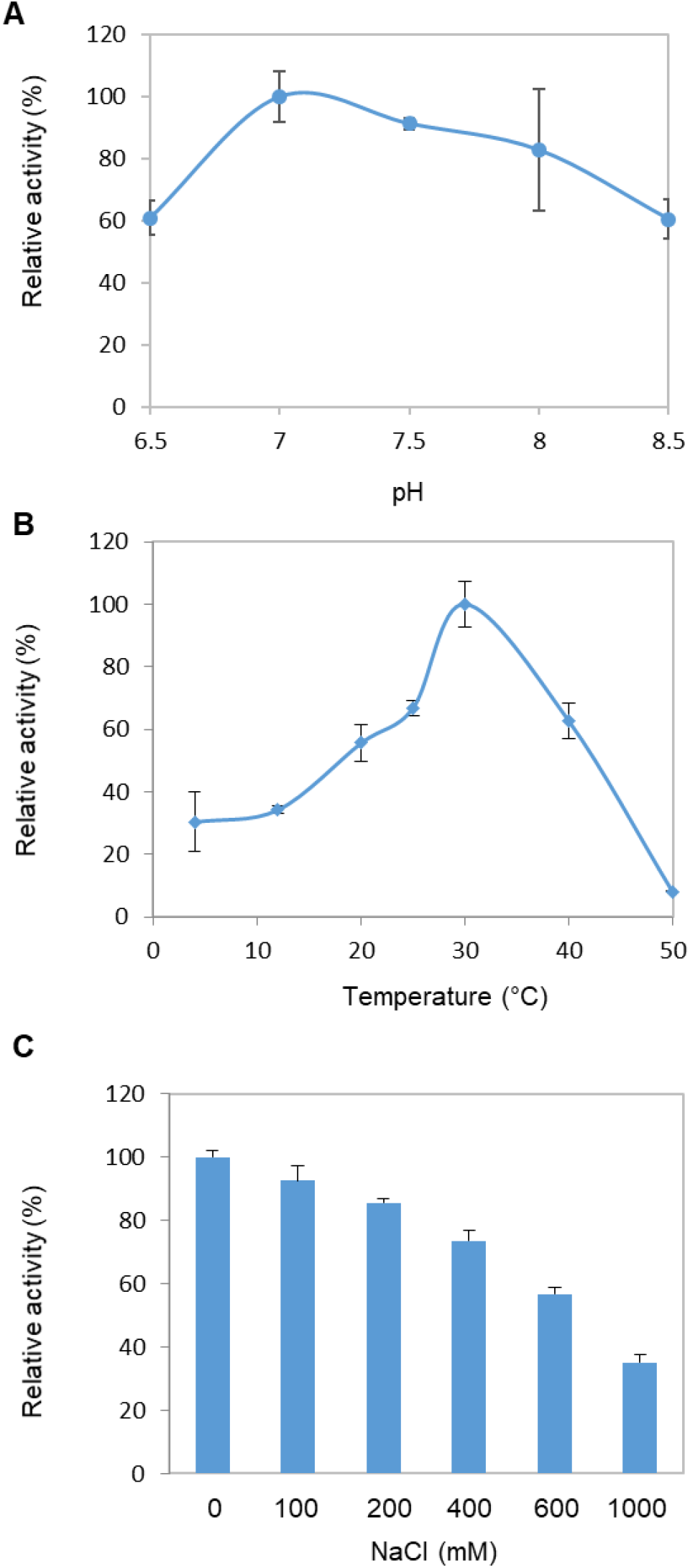
Influence of NaCl concentration (A), temperature (B) and pH (C) on mannitol-1-phosphate hydrolysis activity of recombinant EsM1Pase1short.

### 2.3 Evolution and potential localization of M1Pases in brown algae

Phylogenetic analysis of 38 sequences of biochemically characterized and candidate M1Pases identified in six orders of brown algae (Supplementary File S1) revealed that these proteins grouped into two well-supported clusters (Fig. 4). One contained EsM1Pase1 and SjaM1Pase1, while the other included EsM1Pase2 and SjaM1Pase2. This suggested that the last common ancestor before the evolution of the different brown algal lineages contained the two distinct M1Pases. To complete this analysis, two prediction tools, ASAFind and HECTAR, were used to assess potential subcellular localization of brown algal M1Pases. All the sequences found to contain a signal peptide by ASAFind were predicted to be chloroplastic or to contain a signal peptide by HECTAR (Fig. 4). It is worth to mention that all the potential plastidial proteins were in the cluster corresponding to the M1Pase1 (Fig. 4), whereas the other group corresponds to putative cytosolic M1Pases. These predictions suggested that mannitol production might occur both in the chloroplast and in the cytoplasm. In line with this observation, during analysis of genes involved in central metabolism of the unicellular stramenopile *Nannochloropsis oceanica*, it was predicted that both the M1PDH and M1Pase could be chloroplastic (Poliner et al., 2015). However, none of the three M1PDHs identified in the *Ectocarpus* sp. genome were predicted such a subcellular localization, but most of the brown algal M1PDH1 orthologs were predicted by ASAFind to contain a signal peptide or to be localized in the chloroplast (Tonon et al., 2017). It will thus be interesting to establish experimentally the localization of the mannitol biosynthetic genes in brown algae, and especially in the *Ectocarpus* model, to better understand the spatio-temporal organization of this important metabolic pathway.

**Fig. 4.**
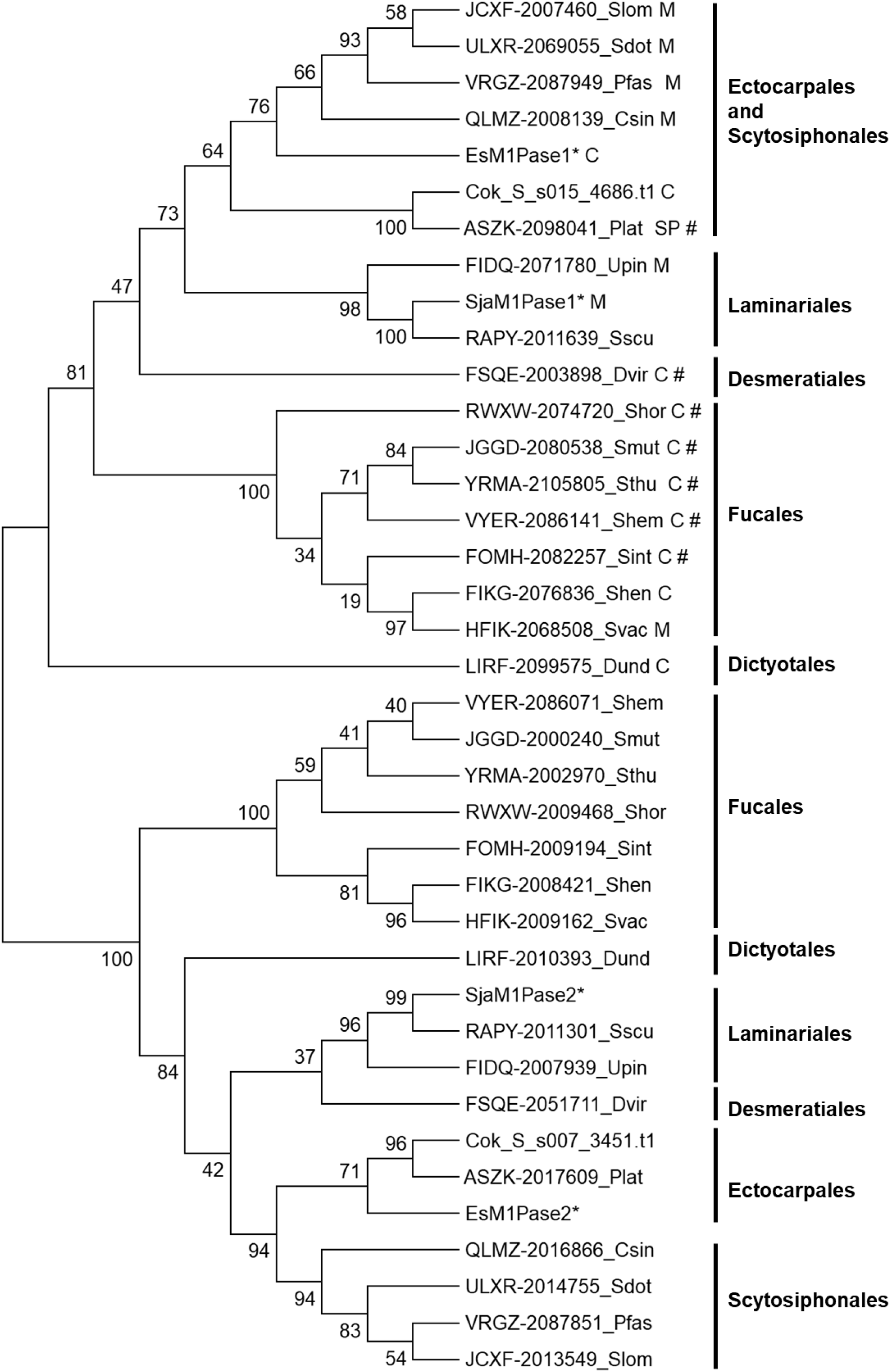
Phylogenetic analysis of brown algal M1Pases and their predicted localization. The evolutionary history was inferred by using the Maximum Likelihood (ML) method based on the JTT matrix-based model. The bootstrap values in the ML analysis are indicated next to the branches (100 replicates). The analysis involved 38 amino acid sequences. All positions with less than 95% site coverage were eliminated. There were 312 positions in the final dataset. The origin of the sequences is indicated by a 4-letter abbreviation at the end of the name of the sequences, except for *Ectocarpus* sp (EsM1Pase1 and EsM1Pase2), *Saccharina japonica* (SjaM1Pase1 and SjaM1Pase2), and *Cladosiphon okamarinus* (Cok_S_s007_3451.t1 and Cok_S_s015_4686.t1) sequences. * indicates recombinant proteins which have been biochemical characterized after expression in *E. coli*. C, M, and SP indicates potential chloroplast localization, mitochondrion localization and presence of a signal peptide predicted by HECTAR (Gschloessl et al. 2008) respectively. # indicates presence of a signal peptide predicted by ASAFind (Gruber et al., 2015).

## 3. Conclusion

The characterization of the second putative M1Pase in the brown alga *Ectocarpus* sp. indicates that both EsM1Pases feature narrow substrate specificity, being active only towards mannitol-1P, and thus are probably specifically involved in mannitol biosynthesis. This in contrast with observation made in the closely related brown alga *S. japonica*. Phylogenetic analysis and prediction of subcellular localization of the two types of M1Pases also suggested that they could have diverse and complementary roles in mannitol metabolism of brown algae. Results presented here also point out that both EsM1Pases have different level of redox-sensitivity, with reducing agent being strictly required for EsM1Pase1 to be active, while significantly increasing EsM1Pase2 activity. To our knowledge, recombinant EsM1Pases enzymes are the second example of redox-sensitive HAD-type phosphatases. This suggests that such regulation mechanisms has been conserved in HAD hydrolases acting on distinct substrates and across different evolutionary lineages, paving the way for further exploration on physiological roles and regulation mechanisms of members of this superfamily of proteins.

## 4. Experimental

### 4.1 Testing of mannitol production by *Ectocarpus* sp. M1Pases in *Escherichia coli*

Recombinant *E. coli* cells expressing native full-length EsM1Pase1 and EsM1Pase2 were obtained as previously described (Groisillier et al., 2014). To improve expression in *E.coli*, gene coding for EsM1Pase1 (Esi0080_0016; UniProt accession number CBJ27643) was codon-optimized (GeneArt Gene Synthesis, Life Technologies, USA), amplified with the forward primer 5’-GGGGGGGGATCCGCGATGAAGCGGACCATACAGG-3’ (*Bam*HI site in italic) and the reverse primer 5’-CCCCCCGAATTCTTATTCCCACACCGTCTTCCTGTCC-3’ (EcoRI site in italic), and cloned into the vector pFO4 to construct the plasmid pESM1Pase1opt. Sequence of this plasmid was verified by sequencing before transformation into in *E. coli*.

Mannitol production in *E. coli* BL21 (DE3) cells transformed with plasmid containing full-length native EsM1Pase1, full-length *E. coli* codon-optimized ESM1Pase1, or EsM1Pase2 was assessed in triplicates for each gene. Pre-cultures of recombinant *E. coli* were grown in five ml of M9 medium supplemented with 10 g/L of glucose and 0.1 g/L ampicillin overnight at 37 °C and 200 rpm, and subsequent experiment conducted as previously described (Madsen et al., 2018). Briefly, these pre-cultures were used the next day to start new cultures at OD_600_ 0.1, and incubated until OD_600_ was 0.5. Cultures were then split in twice five ml for each clone, IPTG (1 mM final concentration) added in one of the two tubes, and culture proceeded 20 hours at 25°C and 200 rpm. After this incubation, cells were harvested by centrifugation at 3,500 *g* for 10 min and supernatant frozen at −20 °C. Mannitol in the extracellular media was quantified using a D-Mannitol/L-Arabitol assay kit (K-MANOL, Megazyme) following manufacturer’s instructions.

### 4.2 Production and purification of *EsM1Pase1short*

The presence of a chloroplast transit peptide of 39 aa at the N terminus of EsM1Pase1 was predicted using HECTAR v1.3 (http://webtools.sb-roscoff.fr; Gschloessl et al., 2008), and the N-terminal boundary was refined by Hydrophobic Cluster Analysis (HCA) (Lemesle-Varloot et al., 1990). Based on this, the *EsM1Pase1shor*t gene corresponding to aa 40 to 405 was amplified with the forward primer 5’-GGGGGG*GGATCC*ACCGCAGCACATGTTAGCGCAG −3’ (*Bam*HI restriction site in italic) and the reverse primer 5’-CCCCCC*GAATTC*TTATTCCCACACGGTTTTGCGATCCA −3’ (*Eco*RI restriction site in italic). This PCR fragment was cloned into the vector pFO4 (adding a six histidine tag at the N-terminus of the recombinant protein) to produce the plasmid pEsM1Pase1short. The integrity of the sequence was verified by sequencing.

*E. coli* strain BL21 (DE3) (Novagen(R)) was transformed with pEsM1Pase1short. To induce production of recombinant protein, a double induction medium was used. For this, transformed *E. coli* were grown in 500 ml of LB containing 0.5% glucose and 100 mg/ml ampicillin at 37°C, and shaked at 180 rpm until OD_600_ reached 1.2-1.5. Protein expression was then induced by adding 500 ml of LB, previously stored at 4°C, 50 ml of lactose 12% (w/v), 20 ml of HEPES 1 M, and IPTG at 0.1 mM final concentration. Cultures were further incubated for 20 hr at 20°C and 180 rpm.

EsM1Pase1short protein was purified using previously described method (Bonin et al., 2015). The procedure included a Ni^2+^ affinity chromatography step, and a size exclusion chromatography separation, after which some fractions were analysed by sodium dodecyl sulfate polyacrylamide gel electrophoresis (SDS-PAGE) using 12% Criterion precast Bis-Tris gels (BioRad). Protein concentration was measured at 280 nm using a Nanodrop 2000 Spectrophotometer (Thermofisher). A molar extinction coefficient of 28,670 M^−1^ cm^−1^, and a molecular weight of 41 kDa, both calculated for the EsM1Pase1short protein sequence at https://web.expasy.org/cgi-bin/protparam, were considered to determine the concentration of the purified enzyme in the fractions of interest.

### 4.3 Biochemical characterization of EsM1Pase1short

If not stated otherwise, all assays were carried out in technical triplicate at 30 °C in 80-μl reaction assays. All compounds used were ordered from Sigma-Aldrich (USA). EsM1Pase1short enzymatic activity was assayed as reported previously (Groisillier et al., 2014), with some modifications. The standard reaction mixtures contained 1 mM Mannitol-1P, 100 mM M Tris-HCl (pH 7.5), 5 mM MgCl_2_, and 3 mM final concentration of reducing agent (DTT). Reactions were initiated by adding about 1 μg of purified recombinant EsM1Pase1short. Free phosphate concentrations were determined using the Malachite Green Phosphate Assay Kit following manufacturer’s instructions (BioAssay Systems, USA). For assessing phosphatase substrate specificity, six sugar and polyol phosphoesters were tested at 1 mM final concentration: mannitol-1P, mannose-6P, fructose-1P, fructose-6P, glucose-1P and glucose-6P. The dependence of enzyme activities on pH and temperature was determined using pH ranging from 5.5 to 9.5 in 0.1M Tris-HCl buffer, and temperature from 10°C to 50°C. Influence of NaCl on enzyme activity was tested in presence of final concentrations ranging from 0 to 1 M. Kinetic parameters were determined after measurement of specific activities in presence of different concentrations of M1P (from 0.0625 to 1.25 mM). The EsM1Pase1short activity was calculated as reported previously (Groisillier and Tonon, 2016).

### 4.4 Retrieval of sequences, prediction of peptide signal, determination of potential subcellular localization, and phylogenetic analysis

All brown algal sequences were retrieved from the OneKP project (transcriptomic resource; https://sites.google.com/a/ualberta.ca/onekp/, Johnson et al., 2012), except for *Ectocarpus* sp., *Saccharina japonica* and *Cladosiphon okamarinus* for which sequences were retrieved from corresponding genomes. All the sequences are given in Supplementary data S1. HECTAR v1.3 (Gschloessl et al., 2008), and ASAFind v1.1.5 (https://rocaplab.ocean.washington.edu/tools/asafind/; Gruber et al., 2015) were used to predict signal peptide, and potential localization to plastids, mitochondria, endoplasmic reticulum, and cytoplasm.

The sequences were aligned with Muscle and phylogenetic analysis were performed with Mega 6.0 as previously described (Tonon et al., 2017).

## Acknowledgements

TT was supported by The Leverhulme Trust Research project Grant RPG-2015-102. The funder had no role in study design, data collection and interpretation, or the decision to submit the work for publication. This work has also benefited from the support of the project IDEALG (ANR-10-BTBR-02) “Investissements d’Avenir, Biotechnologies-Bioressources”.

## References

Abdul Khalil, H. P. S., et al. 2017. Seaweed based sustainable films and composites for food and pharmaceutical applications: a review. Renew. Sust. Energ. Rev. 77, 353–362.

Bonin, P., et al. 2015. Molecular and biochemical characterization of mannitol-1-phosphate dehydrogenase from the model brown alga *Ectocarpus* sp. Phytochemistry 117, 509–520.

Chi, S., et al. 2018a. Functional genomics analysis reveals the biosynthesis pathways of important cellular components (alginate and fucoidan) of *Saccharina*. Curr. Genet. 64, 259–273.

Chi, S., et al. 2018b. Characterization of mannitol metabolism genes in *Saccharina* explains its key role in mannitol biosynthesis and evolutionary significance in Laminariales. bioRxiv, https://www.biorxiv.org/content/10.1101/243402v1.article-info.

Cock, J. M., et al. 2010. The *Ectocarpus* genome and the independent evolution of multicellularity in the brown algae. Nature 465, 617–621.

Dittami S. M., et al. 2009. Global expression analysis of the brown alga *Ectocarpus siliculosus* (Phaeophycaea) reveals large-scale reprogramming of the transcriptome in response to abiotic stress. Genome Biol. 10, R66.

Eggert, A., et al. 2006. Biochemical characterization of mannitol metabolism in the unicellular red alga *Dixoniella grisea* (Rhodellophyceae). Eur. J. Phycol. 41, 1–9.

Fischl, R., et al. 2016. The cell-wall active mannuronan C5-epimerases in the model brown alga *Ectocarpus*: From gene context to recombinant protein. Glycobiology 26, 973–983.

Gravot, A., et al. 2010. Diurnal oscillations of metabolite abundances and genomes analysis provide new insights into central metabolic processes of the brown alga *Ectocarpus siliculosus*. New Phytol. 188, 98–110.

Groisillier, A. 2018. Cloning and expression strategies for the post-genomic analysis of brown algae. In book: Protocols for macroalgae research, 453–468.

Groisillier, A., et al. 2014. Mannitol metabolism in brown algae involves a new phosphatase family. J. Exp. Bot. 65, 559–570.

Groisillier, A., Tonon, T. 2016. Determination of Recombinant Mannitol-1-phosphatase Activity from *Ectocarpus* sp. Bio-protocol 6, e1896.

Gruber, A., et al. 2015. Plastid proteome prediction for diatoms and other algae with secondary plastids of the red lineage. Plant J. 81, 519–528.

-Gschloessl, B., et al. 2008. HECTAR: a method to predict subcellular targeting in heterokonts. BMC Bioinformatics 9, 393.

-Ikawa, T., et al. 1972. Enzymes involved in last steps of biosynthesis of mannitol in brown algae. Plant Cell Physiol. 13, 1017–1029.

Inoue, A., et al. 2016. Functional heterologous expression and characterization of mannuronan C5-epimerase from the brown alga *Saccharina japonica*. Algal Res. 16, 282–291.

Inoue, A., Ojima, T. 2019. Functional identification of alginate lyase from the brown alga *Saccharina japonica*. Sci. Rep. 9, article number 4937.

Iwamoto, K., et al. 2001. Purification and characterization of mannitol-1-phosphatase dehydrogenase in the red alga *Caloglossa continua*. Plant Physiol. 133, 893–900.

Johnson, M. T. J. 2012. Evaluating methods for isolating total RNA and predicting the success of sequencing phylogenetically diverse plant transcriptomes. PLoS One 11, e50226.

Kawai, S., Murata, K. 2016. Biofuel production based on carbohydrates from both brown and red macroalgae: recent developments in key biotechnologies. Int. J. Mol. Sci. 17, 145.

Krause-Jensen, D., et al. Sequestration of macroalgal carbon: the elephant in the Blue Carbon room. Biol. Lett. 14, 20180236; http://dx.doi.org/10.1098/rsbl.2018.0236 (2018).

Lemesle-Varloot, L., et al. 1990. Hydrophobic cluster analysis: procedures to derive structural and functional information from 2-D-representation of protein sequences. Biochimie 72, 555–574.

Liberator, P., et al. 1998. Molecular cloning and functional expression of mannitol-1-phosphatase from the apicomplexan parasite *Eimeria tenella*. J. Biol. Chem. 273, 4237–4244.

Madsen, M. A., et al., 2018. Engineering mannitol biosynthesis in *Escherichia coli* and *Synechococcus* sp. PCC 7002 using a green algal fusion protein. ACS Synth. Biol. 12, 2833–2840.

Michel, G., et al. 2010. Central and storage carbon metabolism of the brown alga *Ectocarpus siliculosus*: insights into the origin and evolution of storage carbohydrates in Eukaryotes. New Phytol. 188, 82–97.

Poliner, E., et al. 2015. Transcriptional coordination of physiological responses in *Nannochloropsis oceanica* CCMP1779 under light/dark cycles. Plant J. 83, 1097–113.

Reed, R. H., et al. 1985. The osmotic role of mannitol in the Phaeophyta: an appraisal. Phycologia. 24, 35–47.

Rioux, L-E., Turgeon, S. 2015. Seaweed carbohydrates. in Seaweed sustainability: food and non-food applications book, 141–192.

Rousvoal, S., et al. 2011. Mannitol-1-phosphate dehydrogenase activity in *Ectocarpus siliculosus*, a key role for mannitol synthesis in brown algae. Planta 233, 261–273.

Sand, M., et al. 2015. Mannitol-1-phosphate dehydrogenases/phosphatases: a family of novel bifunctional enzymes for bacterial adaptation to osmotic stress. Environ. Microbiol. 17, 711–719.

Schiener, P., et al. 2015. The seasonal variation in the chemical composition of the kelp species *Laminaria digitata*, *Laminaria hyperborea*, *Saccharina latissima* and *Alaria esculenta*. J. Appl. Phycol. 27, 363–373.

Seifried, A., et al. 2016. Reversible oxidation controls the activity and oligomeric state of the mammalian phosphoglycolate phosphatase AUM. Free Radic. Biol. Med. 97, 75–84.

Sudhakar, K., et al. 2018. An overview of marine macroalgae as bioresource. Renew. Sust. Energ. Rev. 91, 165–179.

Tenhaken, R., et al. 2011. Characterization of GDP-mannose dehydrogenase from the brown alga *Ectocarpus siliculosus* providing the precursor for the alginate polymer. J. Biol. Chem. 286, 16707–16715.

Tonon, T., et al. 2017. Mannitol biosynthesis in algae: more widespread and diverse than previously thought. New Phytol. 213, 1573–1579.

Wei, N., et al. 2013. Marine macroalgae: an untapped resource for producing fuels and chemicals. Trends Biotechnol. 31, 70–77.

Ye, N., et al. 2015. *Saccharina* genomes provide novel insight into kelp biology. Nat Commun. 6, article number 6986.

Zhang, P., et al. 2016. Comparative characterization of two GDP-mannose dehydrogenase genes from *Saccharina japonica* (Laminariales, Phaeophyceae). BMC Plant Biol. 16, 62.

