## Supplemental data for "Characterization of redox sensitive algal mannitol-1-phosphatases of the haloacid dehalogenase superfamily of proteins"

**Fig. S1.** Kinetic analysis activity of EsM1Pase1short against mannitol-1P. The Lineweaver–Burk plot for $K_m$ and $V_m$ calculation was obtained for mannitol-1P concentrations ranging from 0.0625 to 1.25 mM. For each substrate, values represent means ± SD calculated from three reaction assays for one round of purification.

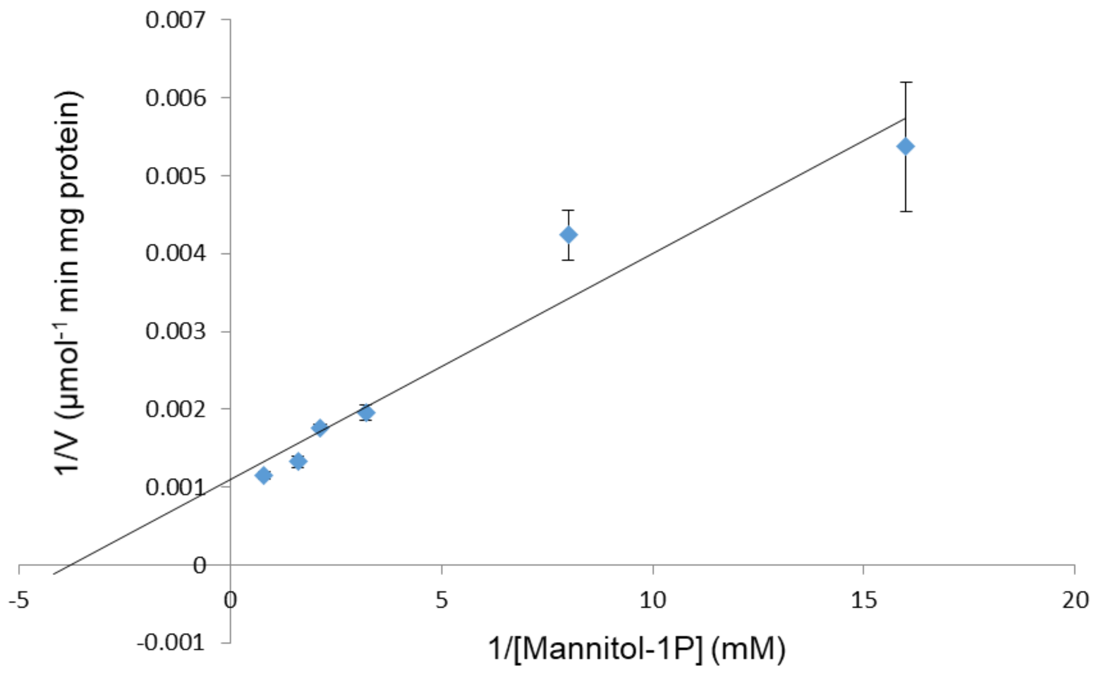

**Fig. S2.** Kinetic analysis activity of EsM1Pase2 against mannitol-1P in presence of 3 mM DTT. The Lineweaver–Burk plot for $K_m$ and $V_m$ calculation was obtained for mannitol-1P concentrations ranging from 0.0625 to 1.25 mM. For each substrate, values represent means ± SD calculated from three reaction assays for one round of purification.

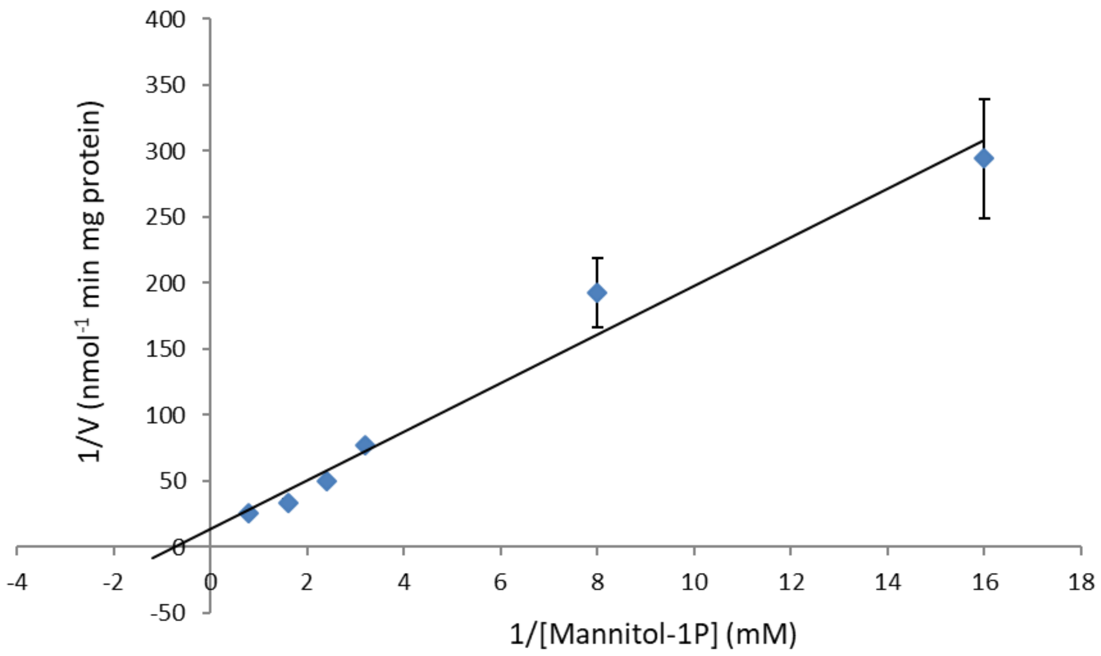

**File S1.** List of brown algal M1Pase sequences used for the phylogenetic analysis, and for the prediction of subcellular localization and presence of peptide signal (Fig. 4).

>EsM1Pase1
MAMKRTIQAAAVLCYHTGATAFLVSTSSSLLSRPLSAATTAAHVSAVRHRRWQQQQQRTARGPTGLSAVEGEEEE
LTDEEKLRRSRLAEDFGANVEVDPKRVAVLFDFDGTIGDTETPAMEVAYWELAPYFPEAAKGGELTDLTEYVRNNA
GKAFEFMLEVVEEDRKAQGLPDIATARAEGAEDPKIMKVVDEARAKYGLKPLGELRAAGTLKDILTQQKEETVDAL
SVVARPVEGMVDTLDELKARKVPFAIATTSPKPRVPASVHACGLDDYFPADKIHSGESDFDPPRFKPDPSVYLRAA
QYEGVIPPICVAVEDSASGVGSAFAAEMGLIVGYVGASHISEERKTEHAKMLRVRGARVVVDNMRDLIPLVDCFTE
CMAKGEDFCLPIANVIKNMDRKTVWE
>Cok_S_s015_4686.t1
MRFLQAAAVLCYHRGATAFILSSSSSSASSLRRATPTAHVSTAAQQQQQQQQQHQQQRRPATGGATVGLRAAA
ADGEGDAGGLTDEERAIKALLKEDFGANVEVDPKRVAVLFDFDGTIGDTETPAMEVAYWELAPYFPQAAKGGEM
VDMSEFVRNNAGKAFEFMLEVVEEDRKAAGLPDIATARAEAAEDPKIMETVDAARAKYGLKSLGELRAAGALKDIL
TQQKEETVDALSIVARPVDGMVETLDELRARKLPFAIATTSPKPRVPASVHACGLDDYFPADKIHSGESDFDPPRFK
PNPSVYLRAAQYEGAIPPLCVAVEDSASGVGSAYAAEMGLIVGYVGASHISEARKNEHAKMLRVRGARVVVNDM
KDLIPLVDCFTECMAKGEDYCLPIAN
LLRTMKGRVWD
>EsM1Pase2
MEKAKAFDKMSILFDFDGTIGDTETPAMEVAFWELAPYLPDTTPDKLDGLMPEFVRDNAGKAFEFMVEKVDEDR
KAAGLSTISEAFAAKSEPKEMLDAVDPHRKKFGLKTFAELRAPDGGEKENLLIQQKTETVDALSKIAQPCNGVPEVL
AALTAAGVAFCISTTSPKPRVPASITACKLDEYFPADKVHSGESDFDPPRFKPNPSVYLKAAETEGKEPANCIAVEDS
GSGVGSASNAGVGLTVGYVGASHIPDYKKDTHAEMLMSGGRAENGKGAEIVISDMTDLPKVIEFFAGEKIAGKSA
PFDFPEELISSLKQPVWVHSTKA
>Cok_S_s007_3451.t1
MERAKAFDKMSILFDFDGTIGDTETPAMEVAFWELAPYLVNTPPEKLDELMPEFVRDNAGKAFEFMVEKVDEDR
KAAGLGTIAEMFAAKSEPKEMLDAVDPHRTKFGLKTFAELRAPDGGEKENLLIQQKTETVDALSKIAQPCNGVPEV
LAALTAAGVPFCISTTSPKPRVPASITACKLDEYFPADKVHSGESDFDPPRFKPNPSVYLKAAETEGKDPVNCIAVEDS
GSGVGSASNAGVGLIVGYVGASHIPEFKKDTHAEMLMAGGRAENGKGAEIVISDMTDLLKVIDFFAGEKTAGKSA
PFEFPADLISSLKKPVWVHGAKA
>SjaM1Pase2
MEQATANKDISILFDFDGTIGDTETPAMEVAFWELAPYLPDTTPDKLDNLMPEFVRDNAGKAFEFMVETVDEERK
AAGMDSVEEMFAAKSEPQNMLDAVDPHRKKFGLKSFAELRAPEGGEAATLLIQQKTETVDALSKIAQPCNGVREV
LAALTAATVPFCISTTSPKPRVPASITACGLDEYFPPDKVHSGESDFDPPRFKPNPSVYLKAAETEGKEPVNCIAVEDS
GSGVGSASNAGVGLTVGYVGASHIPEYKKDTHAEMLMAGGRAENGKGAEIVISDMKDLLKIIDFFAGAKTAGKLA
PFDFPTAMVASMQQPV
WVHGKKA
>SjaM1Pase1
MRRTFQAAAVLCYHRSATAFVSPSAALPSRTSALRRSSTSHSSWVLPQQQVLNEQREQRH
AATALRAGLDGDGSDAEAAMKKLLQEDFGANVAVDPKRVAVLFDFDGTIGDTETPAMEVA
YWELAPYFPGAATGGELVDMSEYVRNNAGKAFEFMLEVVEADRKEQGLPDIATARAESAE
NPEIMKVVDAERAKYGLKPLGDLRAAGTLKDILTQQKEETVDALSVVARPVEGMVNALDE
LRARKLPFAIATTSPKPRVPASIHACGLDDYFPADKVHSGESDFDPPRFKPDPSVYLKAA
QFEGALPPLCVAVEDSASGVGSAYAAEMGLIVGYVGASHISASRKSEHAKMLRHRGARVV
VDEMKDLITLVDCFAECMAKGEDFCLPIANVINTLDPKRVWG
>QLMZ-2016866_Csin
MSLDKAKAFDKMSILFDFDGTIGDTETPAMEVAFWELAPYLPETTPDKLDGLMPEFVRDN
AGKAFEFMVEKVEEDRKAAGMCTIEEAFKAKSEPKEMLDAVDPHRKKFGLKTFAELRAEG
GGEKENLLIQQKTETVEALSKIAQPCNGVPEVLAALTAAGVAFCISTTSPKPRVPASXFP
ADKVHSGESDFDPPRFKPNPSVYLKAAETEGKEPVNCIAVEDSGSGVGSASNAGVGLTVG

YVGASHIPDYKKDTHAEMLMAGGRAENGKGAEIVISDMSDLLKVIDFFAGEKTAGKSAPF
EFPPDLVASMKQPVWVHGKKAQVEASA
>QLMZ-2008139_Csin
MVMKRTMQAAAVLCYHRGATAFVLSSSSASLLRRAPSTAGSTAAHVSASHSQQQRRRSLP
ARAAAARVTALGAGAEGGEAQEDGLTDEERAMKALLKEDFGANVEVDPKRVAVLFDFDGT
IGDTETPAMEVAYWELAPYFPEAAKGGKLTDLTEYVRNNAGKAFEFMLEVVEEDRKAAGL
PDIATARADGAEDPEIMKVVDEARAKYGLKPLGELRAAGTLKDILTQQKEETVEALSVVA
RPVEGMVNTLDELKARKLPFAIATTSPKPRVPASVHACGLDDYFPADKIHSGESDFDPPR
FKPDPSVYLRAAQYEGALPPLCVAVEDSASGVGSAYAAEMGLIVGYVGASHISEARKNEH
AKMLRVRGARVVVNDMKDLIPLVDCFTECMAKGEDFCLPIANLIQTMDKKRVWE
>FSQE-2051711_Dvir
MEKAAANKDISILFDFDGTVGDTETPAMEVAFWELGPYLANTTPDKLDSLMPEFVRDNAG
KAFEFMVETANEDRKAAGLCSVEEMFAAKSEPQDMLDAIDPHRKKFGLKTFAELRAPGGG
ERENLLVQQKTETVDALSKIAQPCNGVREVLATLTSSGVPFCISTTSPKPRVPASITACG
LDEYFPADKVHSGESDFDPPRFKPDPSVYLKAAKTEGKDPVNCIAVEDSGSGVGSASNAG
VGLTVGYVGASHIPDYKKDTHAEMLMAGGRAENGKGAEIVISDMTDLLKIIEFFAGEKTA
GKSAPFEFPSDLVSGLKQPVWVHGQKA
>FSQE-2003898_Dvir
MRRSVQAVVLLCSYRGASAFFSTAVPRASSSLRRTSAPTSSPAHSQQQRQSVVSSASSCV
PVAIAARQSQALRAGGGGDDGEAKVVEDFGANVEVDMKRVAVLFDFDGTIGDTETPAMEV
AFWELAPYFPEAATGGKLEDMPEYIRNNAGKAFEFMLEKVEEDRKTAGLSDIATVRAEAG
ENAEIMKVVDENRAKYGLKPLGELRASGDLRDILTQQKVETVDALSVVARPVEGMVNTLD
ELRARKIPFAIATTSPKPRVPASVTACGLDDYFPGDKIHSGESDFDPPRFKPDPSVYLKA
AQFEGAIPPLCVAVEDSASGVGSADAAEMGLIVGYVGASHIGQARKQEHAKMLRTRGARV
VVDDMKDLIPLVECFTECMAKGEDYCAPIAEVISTLDPKRTWG
>LIRF-2010393_Dund
MLEKATANKNLSILFDFDGTIGDTETPAMEVAFWELGPYLVNTPPSKLESLMPEFVRDNA
GKAFEFMVDTTNEERKAAGMVSIEDMFAAKSEPQEMLDAVDPHREKFGLKKFADLRAEGG
GEKENLLVQQKTETVDALSKIAQPCPGVPEVLAALRAAEVPFCIATTSPKPRVPASITAC
GLDEYFPADKVHSGESDFDPPRFKPNPSVYLKAAETEGKEPANCIAVEDSGSGVGSASNA
GVGLIVGYVGASHIPDFKKDTHAEMLMSGAKSENGKGAEIVISDMSDLLKVINFFAGEKT
EGKSPPFDFPEELIESLKKPVWVQGKKA
>LIRF-2099575_Dund
MVVVTPSPSVAMRRSIQVAAFLCTGRVSAFIASNLRVVSTLRQNSRSSSSLFTFSSPHQQ
EHQGGLCATVSSRRTGTIRAATGSDVEGLVTIEDFGSNVPVDKDRVAVLFDFDGTIGDTE
TPAMEVAYWELAPYFPAAAVGKELIDIREYVRNNAGKAFEFMLEKVEADRAAEGLPDIAT
VRAQAAEHPEIMKVVDEKRAEYGLKSLADLRASGELKDILTQQKEETVDALSVVSRPVQG
MEDTLKELNARTLPFAIATTSPKPRVPASVKACGLEGYFPPDKIHSGESDFDPPRFKPDP
SVYLKAAQFEGALPPLCVAVEDSLSGVGSAANAGIGMIVGYVGASHITEAMKNEHAKSLR
VRGARVVIEDMKNLIPLVECFAGCMAKGADYCEPIAKTIAQLPQKGVWE
>VRGZ-2087851_Pfas
MSLDKAKAFDKMSILFDFDGTIGDTETPAMEVAFWELAPYLPDTTPDKLDGLMPEFVRDN
AGKAFEFMVEKVEEDRKAAGMSTIEEAFKAKSEPKEMLDAVDPHRKKFGLKTFAELRAEG
GGEKEDLLIQQKTETVEALSKIAQPCNGVPEVLAALTAAGVAFCISTTSPKPRVPASITA
CKLDEYFPADKVHSGESDFDPPRFKPNPSVYLKAAETEGKEPVNCIAVEDSGSGVGSASN
AGVGLTVGYVGATHIPDFKMDTHAEMLMAGGRADNGKGAEIVISDMTDLLKIIDFFAGEK
TAGKSAPFEFPPDLVSSLKKPVWVHGSKA
>VRGZ-2087949_Pfas
MAMKRTMQAAAVLCYHRGASAFVLSSSPSSSLLLRRASSRTTAVEAAAHVSASSSQQQRR
RLGGVQPLSAEGEGEAQEDGLSDEERALRALLKEDFGANVEVDPKRVAVLFDFDGTIGDT
ETPAMEVAYWELAPYFPEAAKGGKLTDLTEYVRNNAGKAFEFMLEVVEEDRKTQGLPDIA

TARAEGAEDPEIMKVVDEARAKYGLKPLGELRAAGTLKDILTQQKEETVDALSVVARPVE
GMVNALDELKARKLPFAIATTSPKPRVPASVHACGLDDYFPADKIHSGESDFDPPRFKPD
PSVYLRAAQYEGALPPLCVAVEDSASGVGSAYAAEMGLIVGYVGASHISESRKNEHAKML
RTRGARVVVNDMKDLIPLVDCFTECMAKGEDFCLPIANLIRTMDTRVWE
>ASZK-2017609_Plat
MEKAKAFDKMSILFDFDGTIGDTETPAMEVAFWELAPYLVDTPPEKLDELMPEFVRDNAG
KAFEFMVEKVDEDRKAAGLGTIAXKEMLDAVDPHRQKFGLKTFAELRAPDGGEKENLLIQ
QKTETVDALSKIAQPCNGVPEVLAALTAAGVAFCISTTSPKPRVPASITACKLDEYFPAD
KVHSGESDFDPPRFKPNPSVYLKAAETEGKEPVNCIAVEDSGSGVGSASNAGVGLTVGYV
GASHIPDFKKDTHAEMLMAGGRAENGKGAEIVISDMTDLLKVIEFFAGEKTAGKSAPFEF
PADLISSLKKPVWVHGAKA
>ASZK-2098041_Plat
MRFLQAAAVLCYHRGATAFVLSSSSVSVLRRATATAHVSTAAGAAAGQQQQRHAGLQQGQ
RRAVAAGATGLRAAEGEGAGEESGMTDEERAIKALLKEDFGANVEVDPKRVAVLFDFDGT
IGDTETPAMEVAYWELAPYFPEAATGGNMVDMSEFVRNNAGKAFEFMLEVVEEDRKAAGL
PDIATARAEAAEDPKIMETVDAARAKYGLKPLGELRAAGALKDILTQQKEETVDALSVVA
RPVEGMVNTLDELRARKLPFAIATTSPKPRVPASVHACGLDDYFPADKIHSGESDFDPPR
FKPDPSVYLRAAQYEGAIPPLCVAVEDSASGVGSAYAAEMGLIVGYVGASHISEARKNEH
AKMLRVRGARVVVNDMKNLIPLVDCFTECMAKGEDFCLPIANLVRTMEGVWE
>RAPY-2011301_Sscu
MEKATANKDISILFDFDGTIGDTETPAMEVAFWELAPYLPDTTPDKLDNLMPEFVRDNAG
KAFEFMVETVDEERKAAGMDSVEEMFAAKSEPQNMLDAVDPHRKKFGLKSFAELRAPGGG
EAATLLIQQKTETVDALSKIAQPCNGVREVLAALTAATVPFCISTTSPKPRVPASITACG
LDEYFPPDKVHSGESDFDPPRFKPNPSVYLKAAETEGKEPVNCIAVEDSGSGVGSASNAG
VGLTVGYVGASHIPEYKKDTHAEMLMAGGRAENGKGAEIVISDMKDLLKIIDFFAGAKTA
GKTAPFDFSTDMVASMQQPVWVHGKKA
>RAPY-2011639_Sscu
MTPSHAANCRRPHTHTFHSSGTMRRTFQAAAVLCYHRSATAFVSPSAALPSRTAALRRSS
TSRSSWLLPQEQGLNEQRHAASSATALRAGLDGDGSDAEAAMKKLLKEDFGANVAVDPKR
VAVLFDFDGTIGDTETPAMEVAYWELAPYFPGAATGGELVDMSEYVRNNAGKAFEFMLEV
VEADRKEQGLPDIATARAESAENPEIMKVVDAERAKYGLKPLGDLRAAGTLKDILTQQKE
ETVDALSVVARPVEGMVNALDELRARKLPFAIATTSPKPRVPASIHACGLDDYFPADKVH
SGESDFDPPRFKPDPSVYLKAAQFEGALPPLCVAVEDSASGVGSAYAAEMGLIVGYVGAS
HISASRKSEHAKMLRHRGARVVVDEMKDLITLVDCFAECMAKGEDFCLPIANVINTLDPK
RVWG
>VYER-2086071_Shem
MEKASAYKDMSILFDFDGTIGDTETPAMEVAFWELAPYFPNTRPEQLEGLMPEFIRNNAG
KAFEFMVEKVDEDRKSAGLGTVEEAFAAKSEPKEMLDATDPHRTKFGLKTFAELRAEGGG
EAETILIQQKTETVDALSKIAQPCPGVREVLVALREAGIAFCISTTSPKPRVPASITACG
LDEYFPIEKVHSGESDFTPPEFKPSPAVYLKAAKSEGKEPSNCIAVEDSGSGVGSASNAG
VGLIVGYVGASHIKDDRKEAHAKMLMSGEKSKNGKGAEIVISNMKDLLKIIELFAGERTA
GKSSPFVFPKELVDSLSKPMWVHGGDAE
>VYER-2086141_Shem
MKYSSRTMRLAVVALFSQRSATAFIVSSSPHSSYFSALASRATSTVAAVDSVSEASSLRK
AGKQHRNGARALDSMPGSSHGRATSVLCAIPGGFTDAESALKKLMEEDFGANVEVDPKRV
AVLFDFDGTIGDTETPAMEVAYWELAPYFPAAAKGEELQDMSEFIRNNAGKAFEFMLEVC
EEERKAAGLPDIATVRAEAAEDPEIMKIVDENRAKYGLKPLEELRAAGALKDILTQQKEE
TVEALSVVARPTEGMVEALDELKKRNLPFAIATTSPKPRVPASVHACGLDNYFPPEKIHS
GESDFNPPRFKPDPSVYLKAAQFEGVLPPLSVAVEDSASGVGSADAADMGMIVGYVGASH
ITDERKAEHANTLRLRGARVVINNMKDLIPLVDCFNECMSKGEDYCAPIAKVIETLDKDS
VWE

>FIKG-2008421_Shen
MEKASAYKDMSILFDFDGTIGDTETPAMEVAFWELAPYFPNTSPEQLESLMPEFIRNNAG
KAFEFMVEKVNEDRKSAGLGTVEEAFAAKSESKEMLDATDPHRKKFGLKTFAELRAEGGG
EAETILIQQKTETVDALSKIAQPCPGVREVLVALREAGIAFCISTTSPKPRVPASITACG
LDEYFPIEKVHSGESDFTPPEFKPSPAVYLKAAKSEGKEPSNCIAVEDSGSGVGSASNAG
VGLIVGYVGASHIKDDRKEAHAKMLMSGEKSKNGKGAEIVISNMKDLLKIIELFAGERTA
GKSTPFVFPQELVDSLSKPVWVHGGDAE
>FIKG-2076836_Shen
MKYSSRTMRLAAVALCSQRSAMGFIVSSSPHSSSFSALASRATSTVAAVDSISEASSLRQ
AGKQHRNGARTLDSMPGSSHGRATSALCAIPGGFTDAESALKKLMAEDFGANVEVDPKRV
AVLFDFDGTIGDTETPAMEVAYWELAPYFPAAAKGEELQDMSEFIRNNAGKAFEFMLEVC
EEDRKAAGLPDIATVRAEAAEDPEIMKIVDENRAKYGLKPLGELRAAGALKDILTQQKEE
TVEALSVVARPTKGMVEALDELKKRNLPFAIATTSPKPRVPASVHACGLDDYFPPGKIHS
GESDFNPPRFKPDPSVYLRAAQFEGVLPPLSVAVEDSASGVGSADAADMGMIVGYVGASH
ITDERKAEHANTLRLRGARVVINNMKDLIPLVDCFNECMAKGEDYCAPIAKVIETLDKDS
VWE
>RWXW-2009468_Shor
MEKASTYKDMSILFDFDGTIGDTETPAMEVAFWELAPYFPNTGPEQLEGLMPEFIRNNAG
KAFEFMVEKVDEDRKSAGLGTVEEAFAAKSEPKEMLDATDPHRKKFGLKTFAELRAEGGG
EAETILIQQKTETVDALSKIAQPCPGVREVLVALREAGIAFCISTTSPKPRVPASITACG
LDEYFPIEKVHSGESDFTPPEFKPSPAVYLKAAKSEGKEPSNCIAVEDSGSGVGSASNAG
VGLIVGYVGASHIKDDRKEAHAKMLMSGEKSKNGKGAEIVISNMKDLLKIIELFAGERTA
GKSTPFVFPKELVDSLSKPVWVHGGDAE
>RWXW-2074720_Shor
MKYSSRTMRLAVVALFSQRSATAFIVSSSPHSSSFSALASRAMSTVAAVDSVSEASSLRQ
AGKQYRNEARALDLMPGSSHGRTTSVLCAIPGGFSDAESALKKLMAEDFGANVEVDPKRV
AVLFDFDGTIGDTETPAMEVAYWELAPYFPAAAKGEELQDMSEFIRNNAGKAFEFMLEVC
EEDRKAAGLPDIATVRAEAAEDPEIMKIVDENRAKYGLKPLGELRAAGELKDILTQQKEE
TVEALSVVARPTKGMVEALDELKKRNLPFAIATTSPKPRVPASVHACGLDDYFPPGKIHS
GESDFNPPRFKPDPSVYLKAAQFEGVLPPLSIAVEDSASGVGSADAADMGMIVGYVGASH
ITDERKAEHANTLRLRGARVVINNMKDLIPLVDCFNECMAKGEDYCAPIAKVIDTLDRDS
VWE
>FOMH-2009194_Sint
MEKASAYKDMSILFDFDGTIGDTETPAMEVAFWELAPYFPNTSPEQLESLMPEFIRNNAG
KAFEFMVEKVNEDRKSAGLGTVEEAFAAKSESKEMLDATDPHRKKFGLKTFAELRAEGGG
EADTILIQQKTETVDALSKIAQPCPGVREVLVALREAGVAFCISTTSPKPRVPASITACG
LDEYFPIEKVHSGESDFTPPEFKPSPAVYLKAAKSEGKDPSNCIAVEDSGSGVGSASNAG
VGLIVGYVGASHIKDDRKEAHAKMLMSGEKSKNGKGAEIVISNMKDLLKIIELFAGERTA
GKSTPFVFPQELVDSLSKPVWVHGGDAE
>FOMH-2082257_Sint
MKYSPRTMRLALVALFSQRSATGFIVSSSPHSSSFSALASRATSTVAAVDSISEASSLRQ
AGKQHRNGARTLDSMPGSSHGRATSALRVIPGGFTDAESALKKLMAEDFGANVEVDPKRV
AVLFDFDGTIGDTETPAMEVAYWELAPYFPAAAKGEELQDMSEFIRNNAGKAFEFMLEVC
EEDRKAAGLPDIATVRADAAEDPEIMKVVDENRAKYGLKPLGELRAAGALKDILTQQKEE
TVEALSVVARPTNGMVEALDELKKRNLPFAIATTSPKPRVPASVHACGLDDYFPPGKIHS
GESDFNPPRFKPDPSVYLKAAQFEGVLPPLSVAVEDSASGVGSADAADMGMIVGYVGASH
ITDERKAEHANTLRLRGARVVINNMKDLIPLVDCFNECMAKGEDYCAPIAKVIETLDKDS
VWE
>JGGD-2000240_Smut
MEKASAYKDMSILFDFDGTIGDTETPAMEVAFWELAPYFPNTRPEQLEGLMPEFIRNNAG
KAFEFMVEKVDEDRKSAGLGTVEEAFAAKSEPKEMLDATDPHRKKFGLKTFAELRAEGGG

EAETILIQQKTETVDALSKIAQPCPGVREVLVALREAGIAFCISTTSPKPRVPASITACG
LDEYFPIEKVHSGESDFTPPEFKPSPAVYLKAAESEGKEPSNCIAVEDSGSGVGSASNAG
VGLIVGYVGASHIKDDRKEAHAKMLMSGEKSKNGKGAEIVISNMKDLLKIIELFAGERTA
GKSSPFVFPKELVDSLSKPVWVHDGDAE
>JGGD-2080538_Smut
MKYSSRTMRLAVVALFSQRSATAFIVSSSPHSSYLSALASRATSTVAAVDSVSEASSLRQ
AGKQHRNGARTLDSMPGSSHGRAASVLCAIPGGFTDAETALKKLMEEDFGANVEVDPKRV
AVLFDFDGTIGDTETPAMEVAYWELAPYFPAAAKGEELQDMSEFIRNNAGKAFEFMLEVC
EEDRKAAGLPDIAAVRAQAAEDPEIMKIVDENRAKYGLKSLGELRAAGALKDILTQQKEE
TVEALSVVARPTKGMVEALDELKKRNLPFAIATTSPKPRVPASVHACGLDDYFPPEKIHS
GESDFNPPRFKPDPSVYLKAAQFEGVLPPLSVAVEDSASGVGSADAADMGMIVGYVGASH
ITDERKAEHANTLRLRGARVVINNMKDLIPLVDCFNECMSKGEDYCAPIAKVIETLDKDS
VWE
>YRMA-2002970_Sthu
MEKASAYKDMSILFDFDGTIGDTETPAMEVAFWELAPYFPNTRPEQLEGLMPEFIRNNAG
KAFEFMVEKVDEDRKSAGLGTVEEAFAAKSEPKEMLDATDPHRKKFGLKTFAELRAEGGG
EAETILIQQKTETVDALSKIAQPCPGVREVLVALREAGIAFCISTTSPKPRVPASITACG
LDEYFPIEKVHSGESDFTPPEFKPSPAVYLKAAKSEGKEPSNCIAVEDSGSGVGSASNAG
VGLIVGYVGASHIKDDRKEAHAKMLMSGEKSKNGKGAEIVISNMNDLLKIIELFAGERTA
GKSTPFVFPKELVDSLSKPVWVHSGDAE
>YRMA-2105805_Sthu
MKYSSRTMRLAVVALFSQRSATAFIVSSSPHSSYFSALASRATSTVAAVDSVSEASSLRQ
AGKQNRNGARALDLMSGSSHGRATSALCAIPGGFTDAESALKKLMAEDFGANVEVDPKRV
AVLFDFDGTIGDTETPAMEVAYWELAPYFPAAAKGEELQDMSEFIRNNAGKAFEFMLEVC
EEDRKAAGLPDIATVRAEAAEDPEIMKIVDENRAKYGLKSLGELRAAGALKDILTQQKEE
TVEALSVVARPTKGMVEALDELKKRNLPFAIATTSPKPRVPASVHACGLDDYFPPEKIHS
GESDFNPPRFKPDPSVYLKAAQFEGVLPPLSVAVEDSASGVGSADAADMGMIVGYVGASH
ITDERKAEHANTLRLRGARVVINNMKDLIPLVDCFNECMSKGEDYCAPIAKVIETLDKDS
VWE
>HFIK-2009162_Svac
MEKASAYKDMSILFDFDGTIGDTETPAMEVAFWELAPYFPNTSPEQLESLMPEFIRNNAG
KAFEFMVEKVNEDRKSAGLGTVEEAFAAKSESKEMLDATDPHRKKFGLKTFAELRAEGGG
EAETILIQQKTETVDALSKIAQPCPGVREVLVALREAGIAFCISTTSPKPRVPASITACG
LDEYFPIEKVHSGESDFTPPEFKPSPAVYLKAAKSEGKEPSNCIAVEDSGSGVGSASNAG
VGLIVGYVGASHIKDDRKEAHAKMLMSGEKSKNGKGAEIVISNMKDLLKIIELFAGERTA
GKSTPFVFPQELVDSLSKPVWVHGGDAE
>HFIK-2068508_Svac
MKYSSRTMRLAAVALCSQRSATGFIVSSSPHSSSFSTLASRATSTVAAVDSISKASSLRQ
AGKQHRNGARALDSMPGSSHGRATSALCAIPGGFTDAESALKKLMAEDFGANVEVDPKRV
AVLFDFDGTIGDTETPAMEVAYWELAPYFPAAAKGEELQDMSEFIRNNAGKAFEFMLEVC
EEDRKAAGLPDIATVRAEAAEDPEIMKIVDENRAKYGLKPLGELRAAGALKDILTQQKEE
TVEALSVVARPTKGMVEALDELKKRNLPFAIATTSPKPRVPASVHACGLDDYFPPGKIHS
GESDFNPPRFKPDPSVYLRAAQFEGVLPPLSVAVEDSASGVGSADAADMGMIVGYVGASH
ITDERKAEHANTLRLRGARVVINNMKDLIPLVDCFNECMAKGEDYCAPIAKVIETLDKDS
VWE
>JCXF-2013549_Slom
MSLDKAKAFDKMSILFDFDGTIGDTETPAMEVAFWELAPYLPDTTPDKLDGLMPEFVRDN
AGKAFEFMVEKVEEDRKAAGMSTIEEAFKAKSEPKEMLDAVDPHRKKFGLKTFAELRAEG
GGEKEDLLIQQKTETVEALSKIAQPCNGVPEVLAALTAAGVAFCISTTSPKPRVPASITA
CKLDEYFPADKVHSGESDFDPPRFKPNPSVYLKAAETEGKEPVNCIAVEDSGSGVGSASN
AGVGLTVGYVGASHIPDFKMDTHAEMLMAGGRAENGKGAEIVISDMTDLLKIIDFFAGEK

TAGKSAPFEFPPDLVSSLKKPVWVHGSKA
>JCXF-2007460_Slom
MAMKRTMQAAAVLCYHRGATAFVLSSSPSSLLRRTSSPTTAAAAAAAAXHVSASSQQQQQ
RCRSSRAGAVGGVGPLGAGAEGEVQEDGLTDEERALRALLKEDFGANVEVDPKRVAVLFD
FDGTIGDTETPAMEVAYWELAPYFPEAAKGGKLTDLTEYVRNNAGKAFEFMLEVVEEDRK
TQGLPDIATARAEGAEDPEVMKVVDEARAKYGLKPLGELRAAGTLKDILTQQKEETVEAL
SVVARPVEGMVNALDELKARKLPFAIATTSPKPRVPASVHACGLDDYFPADKIHSGESDF
DPPRFKPDPSVYLRAAQYEGALPPLCVAVEDSASGVGSAYAAEMGLIVGYVGASHISESR
KNEHAKMLRTRGARVVVNDMKDLIPLVDCFTECMAKGEDFCLPIANLVRTMDTKRVWE
>ULXR-2014755_Sdot
MSLDKAKAFDKMSILFDFDGTIGDTETPAMEVAFWELAPYLPDTSPDKLDGLMPEFVRDN
AGKAFEFMVEKVEEDRKAAGMPTIEEVFKAKSEPKEMLDAVDPHRKKFGLKTFAELRAEG
GGEKENLLIQQKTETVEALSKIAQPCDGVPEVLAALTAAGVPFCISTTSPKPRVPASITA
CKLDEYFPADKVHSGESDFDPPRFKPNPSVYLKAAETEGKDPVNCIAVEDSGSGVGSASN
AGVGLTVGYVGATHIPDWKMDTHAEMLMAGGRAENGKGAEIVISDITDLLKIIDFFAGEK
TAGKSAPFEFPPDLVSSLKKPVWVHGSKA
>ULXR-2069055_Sdot
MAMKRTMQAAAVLCYHRGATAFVLSSSPSSLLRRTSSPATAAAPAAAHVSASSQQQQRRR
SSPAGAVGGVSPLGAGAEGEAQEDGLTDEERALRALLKEDFGANVEVDPKRVAVLFDFDG
TIGDTETPAMEVAYWELAPYFPEAAKGGKLTDLTEYVRNNAGKAFEFMLEVVEEDRKAQG
LPDIATARAEGAEDPEVMKVVDEARAKYGLKPIGELRAAGTLKDILTQQKEETVEALSVV
ARPVEGMVNALDELKARKLPFAIATTSPKPRVPASVHACGLDDYFPADKIHSGESDFDPP
RFKPDPSVYLRAAQYEGALPPLCVAVEDSASGVGSAYAAEMGLIVGYVGASHISESRKNE
HAKMLRTRGARVVVNDMKDLIPLVDCFTECMAKGEDFCLPIANLIRTMDTKRVWE
>FIDQ-2007939_Upin
MEKASANKNISILFDFDGTIGDTETPAMEVAFWELAPYLPDTTPDKLDALMPEFVRDNAG
KAFEFMVDTVDEERKAKGMDSIQESFAAKSEPQNMLDAVDPHRKKFGLKTFAELRAPDGG
ELETLLVQQKTETVDALSKIAQPCNGVRDVLAALTAAGVAFCISTTSPKPRVPASITACG
LDEYFPPDKVHSGESDFDPPRFKPNPSVYLKAAETEGKEPVNCIAVEDSGSGVGSASNAG
VGLTVGYVGASHIPEYKKDTHAEMLMAGGRAENGKGAEIVISDMKDLLKIIDFFAGEKTA
GKTAPFDFPTDMVASMQQPVWVHGKKA
>FIDQ-2071780_Upin
MRRTFQAAAVLCYHRSATAFVSPSASLPSGVTTQRRASTSRSFWQQGSLNEQRHDSSSSA
SASRKASSPARSALRASLDSGAEAAMKKLFKEDFGSKATVDPKRVAVLFDFDGTIGDTET
PAMEVAYWELAPYFPAAATGGDLMDMSEYVRNNAGKAFEFMLEVVEADRKEQGLPDIATA
RAEAAEHPEIMKVVDAERAKYGLKPLGDLRAAGTLKDILTQQKEETVDALSVVARPVEGM
VNALDELRARKLPFAIATTSPKPRVPASVHACGLDDYFPADKIHSGESDFDPPRFKPDPS
VYLRAAQYEGALPPLCVAVEDSASGVGSAFAAEMGLIVGYVGASHISAARKSEHAKMLRH
RGARVVVDEMKDLIALVDCFAECMAKGVDFCLPIATVIETLDPKRVWG
